# HGS+ enlarged vesicular compartment serves as site for coronavirus assembly at later infection stage

**DOI:** 10.1101/2025.10.25.684511

**Authors:** Xubing Long, Buyun Tian, Rongrong Chen, Rong Bai, Jinping Yang, Yushuang Fang, Xinyue Fu, Yanhong Xue, Wei Ji, Tao Xu, Zonghong Li

**Affiliations:** The First Affiliated Hospital of Guangzhou Medical University, Guangzhou Medical University, Guangzhou, China; Guangzhou National Laboratory, Guangzhou, China; National Laboratory of Biomacromolecules, CAS Center for Excellence in Biomacromolecules, Institute of Biophysics, Chinese Academy of Sciences, Beijing, China

## Abstract

The precise site of coronavirus assembly remains poorly characterized. In this study, we observed that coronavirus infection induces the host factor hepatocyte growth factor-regulated tyrosine kinase substrate (HGS) to form distinct HGS⁺ enlarged vesicular compartments at later infection stage. Confocal and live-cell super-resolution microscopy showed that viral structural proteins colocalize with these HGS⁺ enlarged vesicular compartments. Correspondingly, APEX2-based electron microscopy (APEX2-EM) and immuno-EM analyses confirmed the presence of assembled virions within these unique HGS⁺ compartments, identifying them as sites of virion assembly. This was further supported by cryogenic correlated light and electron microscopy (cryo-CLEM), which captured ongoing virion formation occurring within HGS⁺ enlarged vesicles. Crucially, whole cell volume EM revealed that HGS deficiency abolishes these vesicular compartments and markedly reduces assembled virions. Lastly, we demonstrated that HGS⁺ vesicular compartments are rearranged from Golgi and endosome/lysosome by coronavirus infection. Together, these findings establish that coronavirus-induced HGS⁺ vesicular compartments function as essential platforms for virion assembly at later infection stage.

## Introduction

The coronavirus life cycle comprises viral entry, replication, assembly, and egress (V’Kovski et al., 2021). Coronaviruses enter host cells via specific receptors, such as angiotensin-converting enzyme 2 (ACE2) for severe acute respiratory syndrome coronavirus 2 (SARS-CoV-2) (Jackson et al., 2022; Yan et al., 2020) and dipeptidyl peptidase 4 (DPP-4) for middle east respiratory syndrome-related coronavirus (MERS-CoV) (Chen et al., 2023; Xiong et al., 2022). After entering host cells, the genomic RNA of coronavirus is translated into two large polyproteins ORF1a and ORF1b. These polyproteins are further degraded by the papain-like protease in non-structural protein 3 (NSP3) and the 3C-like protease in NSP5 into 16 NSPs (NSP1-16) that are involved in genome transcription and replication (Sawicki et al., 2005; V’Kovski et al., 2021). Viral RNA replication occurs within specialized double-membrane vesicles (DMVs), which derive from extensive remodeling of the endoplasmic reticulum (ER) membrane mediated by NSP3 and NSP4 (Chen et al., 2024; Gao et al., 2025; Huang et al., 2024; Ji et al., 2023; Ji et al., 2022; Li et al., 2024; Perry et al., 2025; Yang et al., 2025; Zimmermann et al., 2023). However, the precise sites of coronavirus assembly remain scanty.

Initial electron microscopy (EM) studies of mouse hepatitis virus (MHV) infected cells at 6 to 7 hours post infection (hpi) revealed coronavirus assembly and budding are primarily associated with the ER and Golgi, with viral particles also found in smooth-walled vesicles (Sturman and Holmes, 1983; Tooze et al., 1984). Subsequent studies consistently documented coronavirus assembly and budding within the ER-Golgi intermediate compartment (ERGIC) and the Golgi region during the early infection stage (Klaus et al., 2013; Liang et al., 2025; Saraste and Kuismanen, 1984; Saraste and Marie, 2018; Saraste and Prydz, 2021; Ujike and Taguchi, 2015; Voss et al., 2009). These observations were further corroborated by immuno-EM studies, which showed positive signals for ERGIC and Golgi biomarkers at the sites of virus budding (Ulasli et al., 2010). Since these studies were conducted during early infection (< 8 hpi), when the morphology of cellular organelles remains largely intact, coronavirus assembly and budding are generally thought to take place within the lumen of the ERGIC and Golgi compartments. It should be noted, however, that at this stage viral genome replication predominates, with relatively few virions being produced.

At later stages of infection (e.g., 24 hpi), coronavirus infection leads to fragmentation and reorganization of the Golgi apparatus (Bgatova et al., 2023; Zhou et al., 2017). Under these conditions, no distinct ERGIC or Golgi structures can be identified as sites of virion assembly. Instead, so-called “large virus-containing vesicles” appear to serve as the primary assembly sites (Caldas et al., 2021; Ulasli et al., 2010). Moreover, cryo-electron tomography (ET) studies of SARS-CoV-2-infected cells at 24 hpi have revealed that “single-membrane vesicles” are platforms for viral assembly and budding (Bergner et al., 2022; Mendonca et al., 2021). Volume EM imaging further identified three distinct types of assembly sites in SARS-CoV-2-infected cells: vesicular-tubular compartments, electron-dense, virion-packed vesicles, and multivesicular bodies (Cortese et al., 2020). These structures may either represent alternative secretory pathways or dead-end products of the assembly process. Thus, the precise sites and mechanisms of coronavirus assembly require further clarification.

Our previous study showed that HGS is a critical host factor for pan-coronavirus infection (Long et al., 2025). It directly interacts with viral structural membrane (M) protein and facilitates its trafficking to support virion assembly. As a component of the endosomal sorting complex required for transport (ESCRT), HGS plays a central role in endosomal vesicle budding and the biogenesis of multivesicular bodies (MVBs) (Hurley et al., 2025). It also interacts with numerous cellular proteins and regulates their intracellular trafficking and sorting from the trans-Golgi network to endosomes and lysosomes. In this study, we found that coronavirus infection induces HGS to reorganize into prominent, enlarged vesicular compartments. These structures contain both Golgi and endosomal/lysosomal proteins. Using state-of-the-art light and electron microscopy, we demonstrated that these HGS^+^ enlarged vesicular compartments function as sites of virion assembly during later stages of infection.

## Results and discussion

### Coronavirus infection induces the formation of HGS^+^ enlarged vesicles

While our previous study identified HGS as a critical host factor for pan-coronavirus infection (Long et al., 2025), the cellular response of HGS to coronavirus infection remains unknown. Notably, we observed that MHV infection triggered the formation of HGS**^+^** enlarged vesicles, the number of which increased over the course of infection (Fig. 1A). To determine whether this phenomenon occurs in other coronaviruses, we infected cells with multiple coronaviruses, including SARS-CoV-2, HCoV-229E, HCoV-OC43 and HCoV-NL63. Immunofluorescence (IF) analysis revealed that all coronaviruses tested induced the formation of enlarged HGS-positive vesicles at 20 hpi (Fig. 1B). Collectively, these results showed that the HGS undergoes subcellular localization changes in response to coronavirus infection, suggesting its dynamic role in the host-pathogen interaction.

**Figure 1.**
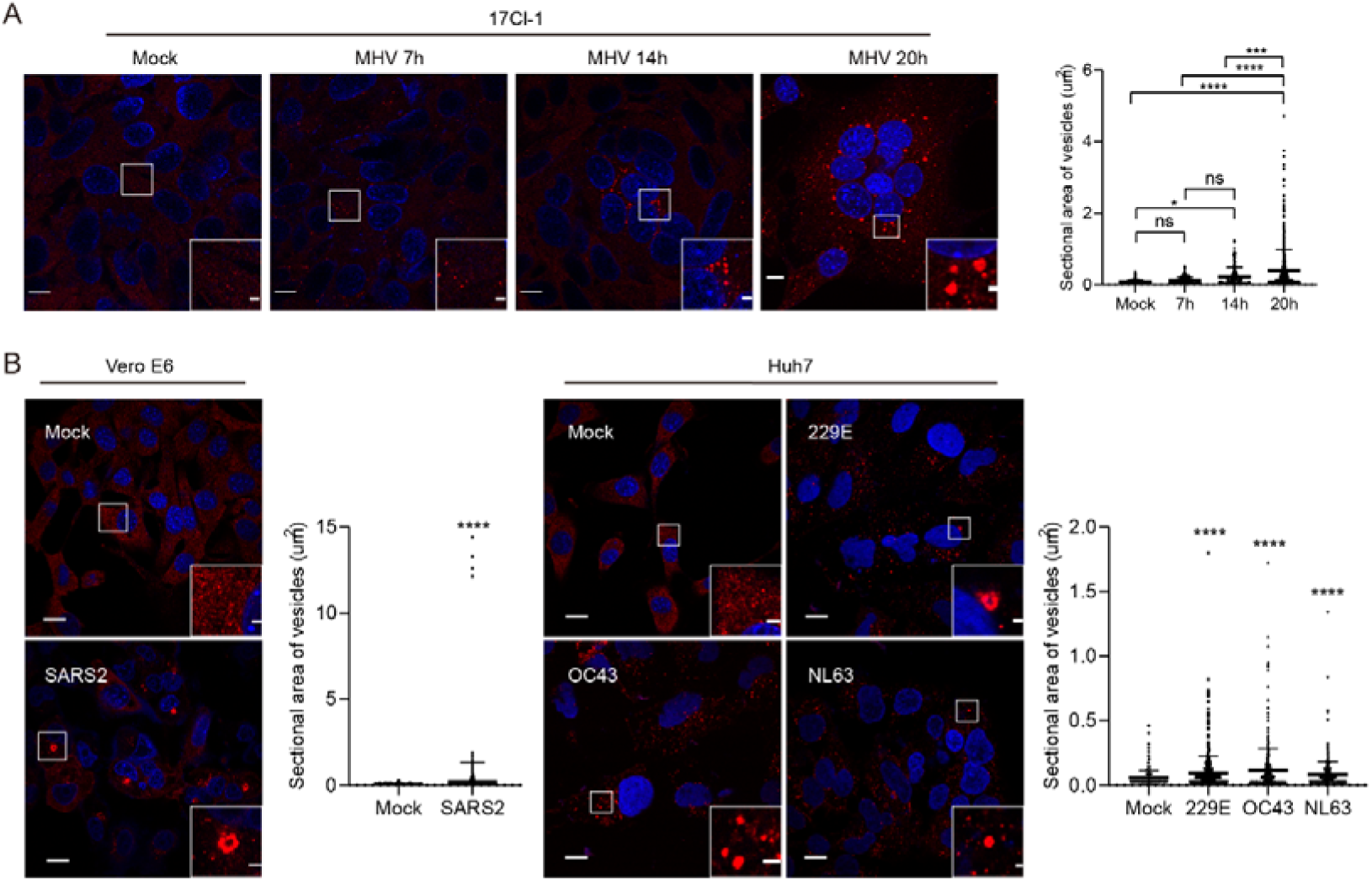
Coronavirus infection induces the formation of HGS-enlarged vesicles. (A) Representative IF analysis of HGS-formed enlarged vesicles (red) in mock-infected or MHV-infected 17Cl-1 cells (MOI = 1, hpi = 7 h, 14 h or 20 h). Nuclear DNA was stained with Hoechst (blue). Scale bar, 10 μm. Scale bar in enlarged view, 1 μm. N = 3 independent biological replications. (B) Representative IF analysis of HGS-formed enlarged vesicles in SARS-CoV-2-infected Vero E6 cells (MOI = 1, hpi = 48 h) and HCoV-229E (MOI = 1, hpi = 48 h), HCoV-NL63 (MOI = 1, hpi = 96 h) or HCoV-OC43 (MOI = 1, hpi = 72 h)-infected Huh7 cells. Scale bar, 10 μm. Scale bar in enlarged view, 1 μm. N = 3 independent biological replications. Data are mean ± SD. Significance for (A-B) was performed with 1-way ANOVA and Tukey’s multiple comparison test. *P ≤ 0.05, **P ≤ 0.005, ***P ≤ 0.0005, ****P ≤ 0.0001, ns, not significant.

### Live-cell imaging indicates that HGS-formed enlarged vesicles are the sites for coronavirus assembly

We next investigated whether HGS⁺ enlarged vesicles are involved in virion assembly. IF assays showed that the viral nucleocapsid (N) protein co-localized with these HGS-positive vesicles in cells infected with MHV, HCoV-229E, or HCoV-OC43. In SARS-CoV-2-infected cells, not only N but also other structural proteins, including the spike (S) and M proteins, were recruited to the HGS⁺ vesicles (Fig. 2A), suggesting these compartments are potential coronavirus assembly sites. Further support came from a SARS-CoV-2 virus-like particle (VLP) assay, in which co-expression of HGS-GFP with M and N proteins resulted in the clear localization of these structural proteins to HGS-defined vesicular structures (Fig. 2B). These findings consistently highlight HGS⁺ enlarged vesicles as functional platforms for coronavirus assembly.

**Figure 2.**
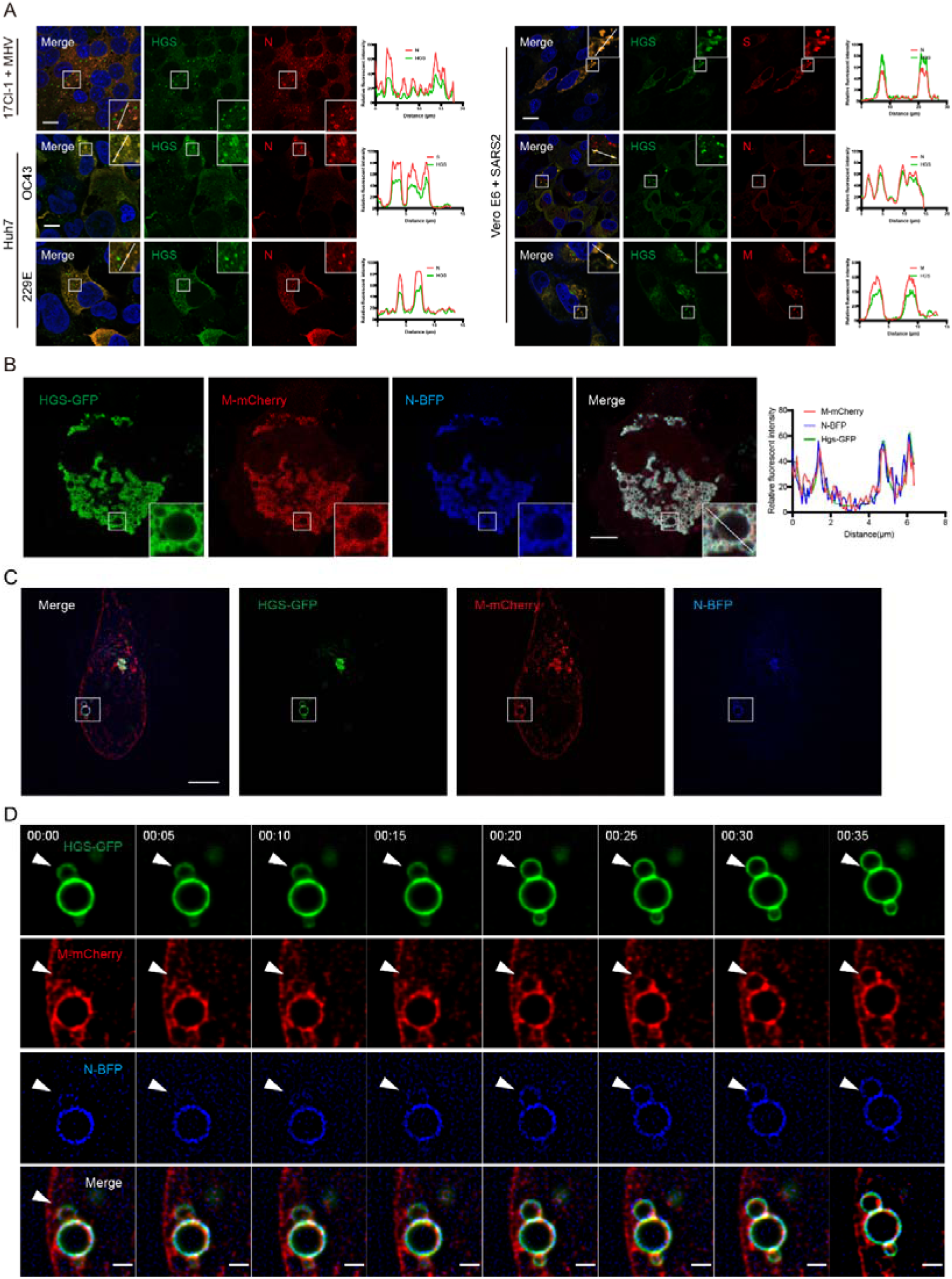
Live-cell imaging illustrates that HGS-formed enlarged vesicles are the sites for coronavirus assembly. (A) Representative IF analysis of viral structural protein co-localization with HGS-formed enlarged vesicles in MHV-infected 17Cl-1 cells, SARS-CoV-2-infected Vero E6 cells, and HCoV-229E- or HCoV-OC43-infected Huh7 cells. Scale bar, 10 μm. N = 3 independent biological replications. (B) Representative IF analysis of HGS-GFP co-localization with SARS-CoV-2 S, M-mCherry and N-BFP. Scale bar, 10 μm. N = 3 independent biological replications. (C-D) HEK293T cells expressing HGS-EGFP, SARS-CoV-2 M-mCherry and N-BFP imaged by Airyscan super-resolution microscopy every 5Ds (C), the time-lapse images of the region outlined by the yellow box in a shown sequentially (D). Arrows indicate sequential recruitment of M and N proteins to HGS-formed vesicles. Scale bar, 500Dnm.

We then performed super-resolution time-lapse imaging of live cells to demonstrate that the HGS-formed enlarged vesicles serve as sites for virion assembly (Fig. 2C and 2D). The results showed that HGS-GFP co-localized with viral structural proteins M-mCherry and N-BFP, representing vesicular structures. Throughout the time-lapse imaging process, we clearly observed that the viral structural proteins M-mCherry and N-BFP were recruited to the HGS^+^ enlarged vesicles (Fig. 2C and 2D). This dynamic visualization provides valuable insight into the molecular interactions occurring during coronavirus assembly within host cells.

### APEX2-EM, immuno-EM and cryo-CLEM illustrate that HGS-formed enlarged vesicular compartments are the sites for coronavirus assembly

To further determine whether these HGS⁺ enlarged vesicles contain virions, we employed APEX2-EM by fusing HGS with APEX2, an engineered peroxidase that serves as an electron-dense tag for EM (Lam et al., 2015). The results indicated that HGS localized to both leaflets of the limiting membranes of the enlarged vesicles (Fig. 3A), confirming the role of HGS in forming these structures. In uninfected cells, the lumens of these vesicles appeared empty (Fig. 3A and 3B). In contrast, in MHV-infected cells, assembled virions were clearly visible within the lumens of HGS-formed enlarged vesicles (Fig. 3C and 3D). Furthermore, using immuno-EM to examine endogenous HGS-labeled structures in infected cells, we confirmed the presence of large assembled virions within vesicles tagged with HGS-specific gold particles (Fig. 3E-H).

**Figure 3.**
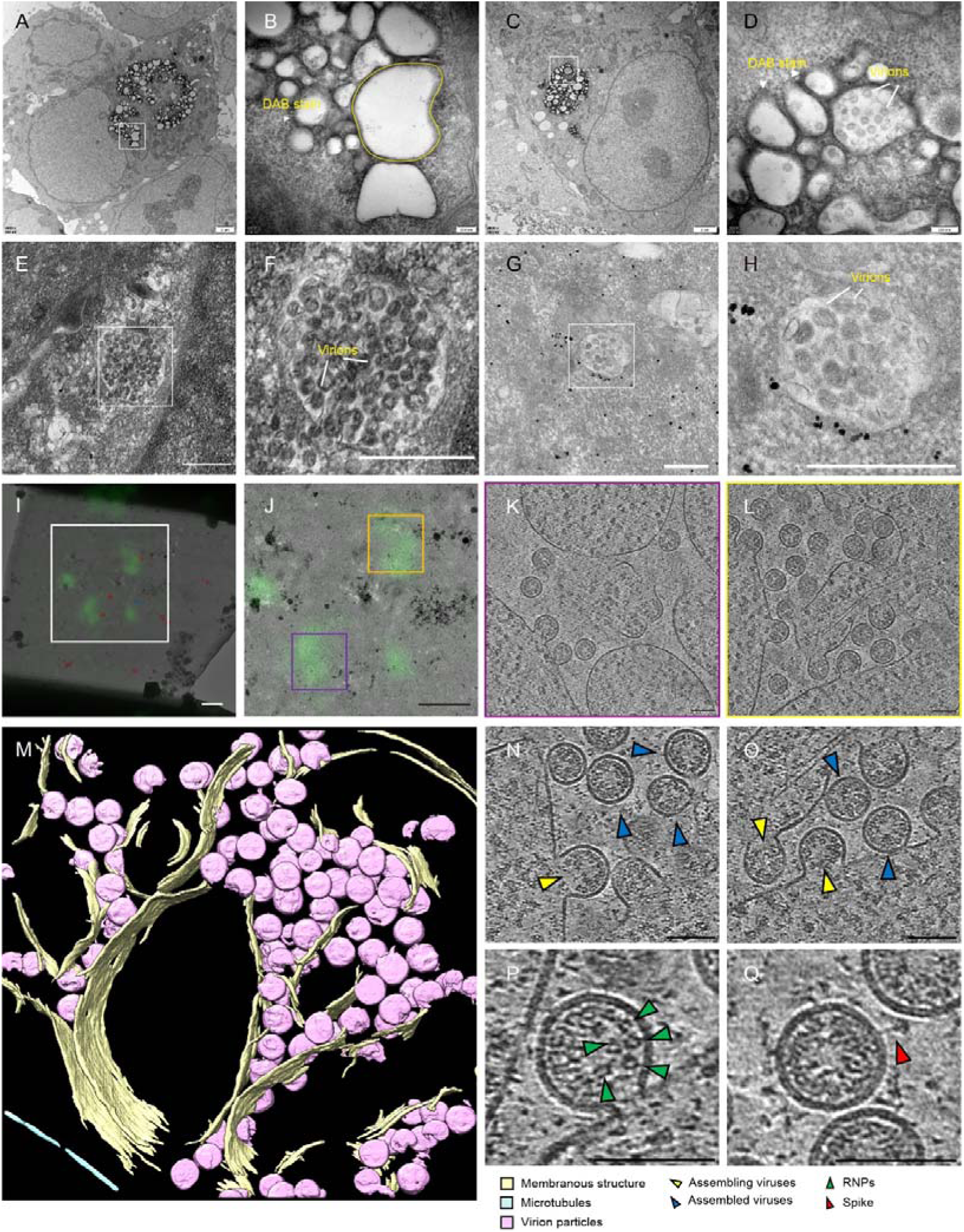
APEX2-EM, immuno-EM and cryo-CLEM illustrate that HGS-formed enlarged vesicles are the sites for coronavirus assembly. (A-D) APEX2-EM of HGS-formed virion-containing vesicles. HEK293T cells transfected with CEACAM1a (MHV receptor) and EGFP-hHGS-APEX2 infected without (A and B) or with (C and D) MHV (MOI = 1, hpi = 20 h). DAB staining localizes HGS to enlarged vesicles containing virions. (B) Enlarged view of white-boxed area in A. (D) Enlarged view of white-boxed area in C. Scale bars: 2 μm in (A) and (C); 200 nm in (B) and (D). (E-H) Immuno-EM showing that HGS is localized on the virion assembling vesicles in MHV-infected 17Cl-1 cells. (E and F) are negative controls (no primary antibody). (F) Enlarged view of white-boxed area in E. (G and H) Immunogold labelling with anti-HGS antibody demonstrate the presence of HGS on the virion assembling vesicles. Gold particles (black dots) indicating the position of HGS molecules. (G) Enlarged view of white-boxed area in H. Scale bars: 500 nm. (I-Q) Cryo-CLEM analysis of HGS-GFP-expressing 17Cl-1 cells infected with MHV (MOI = 1, hpi = 10 h). (I) GFP-HGS fluorescence overlaid on cryo-EM montage. (J) Enlarged view of white-boxed area in I. (K and L) Tomographic slices from boxed area in J, showing viral assembly within HGS-localized vesicles. (M) 3D segmentation of tomogram L. (N-Q) Gallery of MHV virions: assembling virus (yellow arrowheads), assembled viruses (blue arrowheads), RNP density (green arrowheads), MHV spike (red arrowheads). Scale bars: 1 μm (I and J), 200 nm (K and L), 100 nm (N-Q).

To directly demonstrated that the HGS-formed enlarged vesicular compartments are the sites for coronavirus assembly, we performed cryo-CLEM on cryo-focused ion beam milled lamellae in HGS-GFP transfected cells infected with or without MHV. To increase the targeting efficiency, the lamellae were prepared by a cryogenic correlated light, ion and electron microscopy (cryo-CLIEM) technique under the guidance of three-dimensional confocal imaging (Li et al., 2023). In uninfected cells, cryo-ET results revealed multiple enlarged vesicles containing numerous small vesicular cargoes in the HGS-GFP localization (Supplementary Fig. 1A-E). In MHV-infected cells, cryo-ET images displayed abundant virions within the HGS-formed enlarged vesicles (Fig. 3I-Q and Video S1). Notably, well-assembled virions containing structured ribonucleoprotein (RNP) complexes were clearly visible within the vesicular lumens (Fig. 3P and 3Q). Moreover, assembling virions were observed at the limiting membranes of convoluted HGS⁺ vesicular compartments (Fig. 3N and 3O). Spike structures were rarely detected in MHV virions (Fig. 3Q), consistent with previous cryo-ET reports (Bárcena et al., 2009), likely due to the inherent characteristics of MHV. In contrast, cryo-CLEM of WIV1-infected cells revealed virions within HGS⁺ vesicles exhibiting prefusion spike protein crowns (Supplementary Fig. 1F-J). Together, these findings indicate that coronavirus infection induces HGS to generate novel enlarged vesicular organelles that function as central hubs for virion assembly.

### Whole cell volume 3D analysis reveals that HGS deficiency results in the complete loss of virion-assembling vesicular compartments and a dramatic reduction in both the number and size of assembled virions

To obtain a global view of virion assembly sites, we performed focused ion beam-scanning electron microscope (FIB-SEM) analysis of MHV-infected 17Cl-1 cells at a late infection stage (20 hpi, MOI = 1). Full volumes of both wild-type (WT) and *Hgs*-KO infected cells (identified by the presence of DMVs; dimensions 26 × 26 × 10 µm) were acquired. We then employed a bottom-up approach combining semi-automated and automated segmentation using a multicut pipeline to achieve in-depth segmentation of the datasets.

3D reconstruction and visualization of the fully segmented volumes revealed that both WT and *Hgs*-KO infected cells contained multiple distinct nuclei but lacked clearly defined plasma membranes (Video S2 and Video S3), suggesting these cells represent multinucleated giant cells or syncytia. An uninfected cell was also identified within the segmented volume of both conditions (Video S2 and Video S3). Notably, numerous enlarged autophagolysosomes were observed in *Hgs*-KO cells (Supplementary Fig. 2A-C), consistent with previous reports (Coudert et al., 2021; Dauner et al., 2017; Migliano et al., 2024). In both WT and *Hgs*-KO infected cells, endomembrane systems and organelles were markedly altered (Fig 4A and 4B); for instance, mitochondria appeared rounded and had lost most cristae (Supplementary Fig. 2D-I), contrasting with earlier reports of coronavirus-induced mitochondrial elongation (Cortese et al., 2020; Hirabayashi et al., 2025; Shin et al., 2024). We speculate that this discrepancy may reflect differences in infection stages, particularly since previous EM studies showed that the endomembrane system remained relatively normal compared to what was observed here.

**Figure 4.**
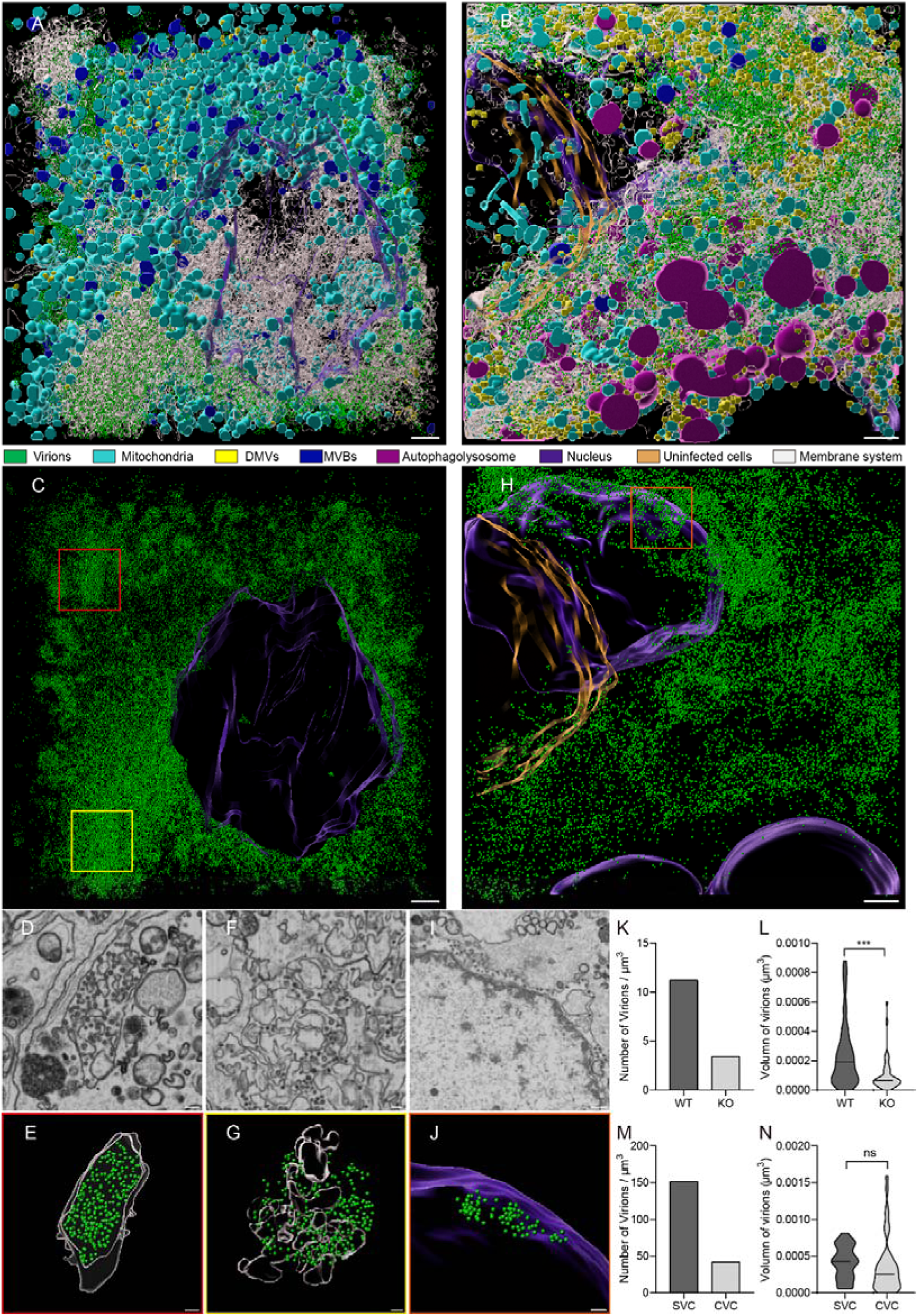
Volume EM reveals that HGS deficiency resulted in the complete loss of virion-assembling vesicular compartments and a dramatic reduction in both the number and size of assembled virions. (A, B) 3D rendering of WT (A) or *Hgs*-KO (B) 17Cl-1 cells infected with MHV (MOI = 1, hpi = 20 h). The color code of subcellular structures is depicted on the bottom. Scale bars, 2 μm. (C, H) 3D rendering of virion and nucleus in WT (A) or *Hgs*-KO (B) 17Cl-1 cells infected with MHV. Scale bars, 2 μm. (D-G) Zoom-in of the area indicated with rectangle in C showing two distinct types of virion assembly events, SVC (D and E) and CVC (F and G), in infected WT cells. Scale bars, 500 nm. (E) same region as in (D) with superimposed rendering of SVC and virion. (G) same region as in (F) with superimposed rendering of CVC and virion. (I-J) Zoom-in of the area indicated with rectangle in H showing virion assembled in perinuclear region of *Hgs*-KO infected cells. Scale bars, 500 nm. (J) same region as in (I) with superimposed rendering of perinuclear region and virion. (K-L) Quantification of virion density (K) and volume of virion (L) in WT and *Hgs*-KO cells. (L) 46 and 47 virions were selected randomly and calculated in WT and *Hgs*-KO cells, respectively. (M-N) Quantification of virion density (M) and volume of virion (N) in SVC and CVC in WT cells. (N) 36 and 43 virions were selected randomly and calculated in SVC and CVC in WT cells, respectively. Data are the mean ± SD. Significance testing for (L and N) was performed with an unpaired t test. *P ≤ 0.05, **P ≤ 0.005, ***P ≤ 0.0005, ****P ≤ 0.0001, ns, no significance.

Interestingly, numerous DMVs were found in close proximity to mitochondria, implying potential functional crosstalk (Supplementary Fig. 2D-I). In WT infected cells, many lysosomes contained electron-dense luminal cargoes, including virions (Supplementary Fig. 3A-B), suggesting a possible cellular defense response to viral infection. Two distinct types of virion assembly events were observed in WT cells: convoluted vesicular compartments (CVCs) and separated vesicular compartments (SVCs) (Fig. 4C-G). Notably, CVCs were found adjacent to structures resembling Golgi stacks (Supplementary Fig. 3C-D), indicating that CVCs may be derived from rearranged Golgi membrane, while SVCs were more likely a fragmentation and redistribution of the SVCs or Golgi cisternae. Strikingly, both types of virion assembly events (CVCs and SVCs) were absent in *Hgs*-KO cells (Video S3). However, a small number of virions were detected in the perinuclear region (Fig. 4H-J), indicating that this zone may serve as an alternative assembly site in the absence of HGS. Quantitative analysis revealed a significantly higher virion density (Fig. 4K) and larger virion volume (Fig. 4L) in WT cells compared to *Hgs*-KO cells. These results suggest despite the presence of an alternative assembly site in *Hgs*-KO cells, the virions produced there are likely structurally or functionally impaired. Moreover, in WT cells, SVCs exhibited higher virion density than CVCs despite comparable virion sizes, indicating a more substantial role for SVCs in virion assembly (Fig. 4M and 4N). In contrast, the number and volume of DMVs were substantially increased in *Hgs*-KO cells compared to WT cells (Supplementary Fig. 2J and 2K), possibly compensating for the loss of conventional virion assembly sites. Collectively, these volume EM data demonstrate that HGS⁺ vesicular compartments constitute the major sites for virion assembly during late stages of coronavirus infection.

### HGS-formed enlarged vesicular compartments incorporate Golgi and endosomal/lysosomal proteins

To explore the composition of the HGS⁺ vesicular compartments, we investigated the protein interactors of HGS under MHV infection condition. The most prominent interacting protein of both mock and MHV-infected conditions identified was STAM, with which HGS forms the heterodimeric ESCRT-0 complex (Fig. 5A). This interaction confirms the specificity of our immunoprecipitation assay. Strikingly, comparative analysis revealed a significant enrichment of Golgi and endosomal/lysosomal proteins within the HGS interactome upon MHV infection (Fig. 5B and 5C), indicating that the HGS-formed enlarged vesicular compartments incorporate components from these organelles during viral infection. To validate this finding, we selected representative markers: a Golgi protein (COG7), an endosomal protein (SNX2), and a lysosomal protein (VPS11). In uninfected cells, their localization with HGS was distinct: COG7 exhibited no colocalization, SNX2 showed partial overlap, and VPS11 completely colocalized with HGS (Fig. 5D), consistent with HGS’s established role as a lysosomal protein (Chin et al., 2001; Eastman et al., 2005; Kanazawa et al., 2003; Kobayashi et al., 2005; Pons et al., 2008; Tamai et al., 2007). However, following coronavirus infection, all selected proteins (COG7, SNX2, and VPS11) demonstrated extensive colocalization with HGS (Fig.5D). This convergence of organellar markers indicates that infection induces the formation of a unique hybrid organelle. The formation of novel organelles was found to be dependent on HGS, as demonstrated by the lack of co-localization between Golgi and lysosomal markers in *Hgs*-KO 17Cl-1 cells (Fig. 5E). All these results suggested that coronavirus infection triggers HGS to form a distinctive enlarged vesicular organelle.

**Figure 5.**
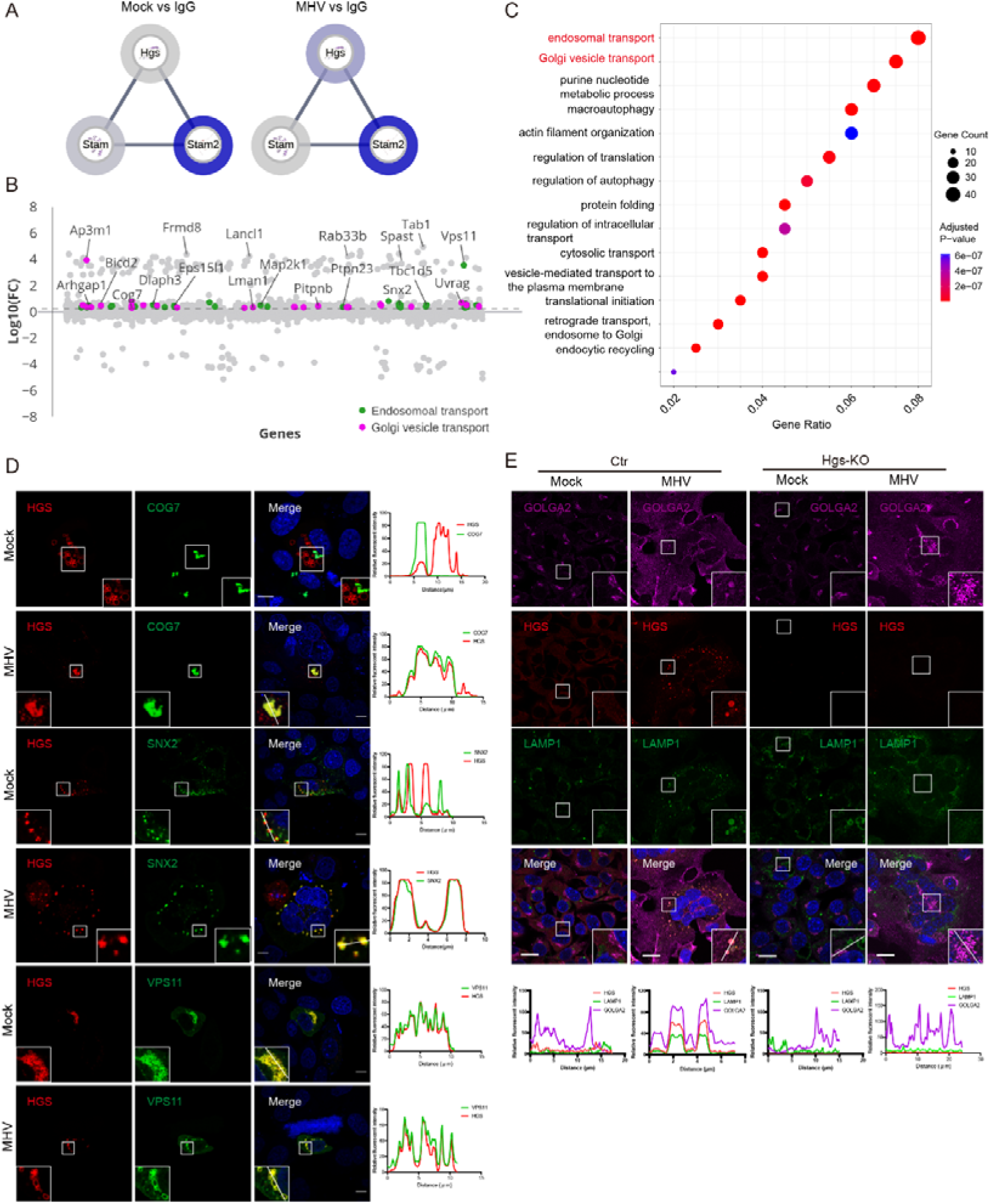
HGS-formed enlarged vesicular compartments incorporate trans-Golgi and lysosome proteins. (A) Protein-protein interaction (PPI) map depicting the HGS-host interactions in mock or MHV-infected 17Cl-1 cells. The halo color indicates the strength of the PPI. (B) Comparative analysis of the HGS-interactome between WT cells and MHV-infected cells. Proteins were colored by Cellular Component. Dotted line indicates fold change (FC) = 2. (C) Gene ontology (GO) analysis of the enriched HGS-interactome in MHV-infected cells. (D) Representative IF analysis of HGS-mCherry co-localization with COG7 (Golgi), SNX2 (endosome) or VPS11 (lysosome) in 17Cl-1 cells infected with or without MHV (MOI = 1, hpi = 20 h). Scale bar, 10 μm. N = 3 independent biological replications. (E) Representative IF analysis of HGS co-localization with GOLGA2 (Golgi) and LAMP1 (lysosome) in WT or *Hgs*-KO 17Cl-1 cells infected with or without MHV. Scale bar, 10 μm. N = 3 independent biological replications.

In conclusion, our study demonstrates that HGS⁺ enlarged vesicular compartments function as essential platforms for virion assembly during the late stages of infection. While the virus utilizes the ERGIC and Golgi apparatus for virion formation in the early infection phase, these organelles become insufficient to support the increasing demand for large-scale virion production as infection progresses. Consequently, the virus induces a reorganization of the host endomembrane system to establish specialized virion assembly sites. In this study, using whole-cell volume EM, we identified two major distinct virion assembly sites at later infection stages, CVCs and SVCs. CVCs are membranes in close proximity to Golgi stacks, resembling the “vesicular-tubular compartments” described in other study (Cortese et al., 2020). SVCs are analogous to “large virus-containing vesicles” or the “single membrane vesicles” reported in SARS-CoV-2-infected cells (Bergner et al., 2022). Although defining the relationship between these compartments remains challenging, we speculate that SVCs likely originate from CVCs.

Although both our study and previous studies have identified large vesicular compartments as virion assembly compartments (Eymieux et al., 2021; Klein et al., 2020; Long et al., 2025; Saraste and Prydz, 2021), the mechanisms underlying their formation and their specific characteristics remain poorly understood. In this study, we reveal that coronavirus infection triggers the formation of enlarged HGS⁺ vesicular compartments and, employing state-of-the-art imaging, confirm that these structures are active virion assembly sites.

Critically, whole-cell volume EM confirms that HGS deficiency abolishes the formation of these virion assembly sites and significantly reduces virion production. Furthermore, we characterized the composition of the HGS⁺ compartments induced by infection and found that they incorporate both Golgi and endosomal/lysosomal proteins, suggesting that they originate from a virus-driven reorganization of these organelles.

Building on our earlier work identifying HGS as a critical host factor that interacts directly with the viral M protein to facilitate virion formation (Long et al., 2025). we propose a model wherein HGS recruits viral structural proteins, primarily via interaction with M, to these compartments to nucleate virion assembly. Finally, in line with previous reports that coronaviruses traffic to lysosomes and egress via Arl8b-dependent lysosomal exocytosis (Yin and Yang, 2024), we suggest that virions are assembled within these HGS⁺ compartments and are subsequently released through the same compartments, thereby utilizing a lysosome-mediated exocytic pathway for efficient viral egress.

## Acknowledgments

The TEM and cryo-CLIEM studies were performed at the Center for Biological Imaging (CBI), Institute of Biophysics, Chinese Academy of Sciences. We thank Xixia Li, Zhongshuang Lv, Xueke Tan and Can Peng at CBI for their assistance in EM sample preparations. The FIB-SEM and Immuno-EM were performed at the Advanced Bioimaging Core Facility (ABCF), Guangzhou National Laboratory, and we would be grateful to Landi Sun, Jiale Zhang, Guosheng Cui and Pei Wang at ABCF for their assistance in EM sample preparations and electron microscopy imaging. Additionally, we are grateful to Ying Li and Qing Zhang from the Proteomics and Metabolomics Core Facility of Guangzhou National Laboratory for their help with MS sample preparation and analysis.

This work was supported by the National Natural Science Foundation of China (92469107 to Z.L.), Major Project of Guangzhou National Laboratory (GZNL2024A01011 to Y.X.; SPRG22-002 to Z.L.), National Key R&D Program of China (2024YFA1307400 to Y.X.), Guangdong Province High-level Talent Youth Project (2021QN02Y939 to Z.L.).

## Author contributions

X.L. performed most viral experiments and IF. B.T. carried out the TEM experiments. R.C. performed the IF and AP-MS data analysis. J.Y. performed the live-cell imaging. R.B. helped with SARS-CoV-2 infection. Y.F. and X.F. helped with FIB/SEM data analysis. Z.L. initiated the study, directed the research and wrote the manuscript. All the authors discussed the data and reviewed the manuscript.

## Methods

### Cell culture and virus

HEK293T, 17Cl-1, LLC-MK2, Vero E6, Huh7 and Huh7.5.1 cells were cultured in Dulbecco’s Modified Eagle’s Medium (DMEM). HRT-18 cells were cultured in RPMI 1640 medium. All cells were cultured in indicated medium supplied with 10% fetal bovine serum (FBS) and 1% antibiotics at 37 °C with 5% CO_2_. A list of cell lines used in this study is provided in the Supplementary Table S1.

MHV virus was a gift from Prof. Hongyu Deng (Institute of Biophysics, Chinese Academy of Sciences). HCoV-OC43 (ATCC VR-1558), HCoV-229E (ATCC VR-740), and HCoV-NL63 (ATCC NR-470) were gifted from Prof. Jincun Zhao (Guangzhou National Laboratory). WIV1 was provided by Prof. Peng Zhou (Guangzhou National Laboratory). SARS-CoV-2 (Genebank accession no. MT123290.1) was a clinical isolate obtained from the First Affiliated Hospital of Guangzhou Medical University. MHV, HCoV-OC43, HCoV-229E, HCoV-NL63 and SARS-CoV-2 were propagated in 17Cl-1, HRT-18, Huh-7, LLC-MK2, and Vero E6 cells, respectively. Viral stocks were stored at −80 °C. SARS-CoV-2 infection was conducted in Guangzhou Customs District Technology Center BSL-3 Laboratory and Guangzhou National Laboratory BSL-3 Laboratory, while other infections were carried out in BSL-2 laboratory of Guangzhou National Laboratory.

### Virus infection

For MHV infection, viruses were added into the 17Cl-1 culture medium at MOI = 1, 2 hpi, culture medium was replaced by fresh complete medium after 2 times wash of PBS, and infected cells were maintained at 37 °C with 5% CO_2_ to collect infected cell lysate for afterwards analysis. For HCoV-OC43, HCoV-229E and HCoV-NL63 infection in Huh7 or Vero E6, after adsorption at 34 °C for 2 h (MOI = 1), the virus inoculum was removed and cells were washed with PBS. Fresh medium was replaced and cells were maintained at 34 °C with 5% CO_2_ to collect infected cell lysate. For SARS-CoV-2 infection in Huh7, Huh7.5.1 or Vero E6, after adsorption at 37 °C for 2 h (MOI = 0.5), the virus inoculum was removed and cells were washed with PBS. Fresh medium was replaced and cells were maintained at 37 °C with 5% CO_2_ to collect infected cell lysate.

### Plasmids

The cDNA fragments of various organelle biomarkers such as LAMP1, SEC61B, TGN38 were amplified by Phanta MaxSuper-Fidelity DNA polymerase (Cat. No. P505-d2) and cloned into the piRFP-N1 vector by ClonExpress II One Step Cloning Kit (Cat. No. C112-02). Homo sapiens HGS, M (SARS-CoV-2), N (SARS-CoV-2) were cloned into the pEGFP-N1, pmCherry-N1 and pBFP-N1 vector, respectively. Sequences of all the inserted DNAs were confirmed by sequencing (Rui Biotech, China).

### Generation of KO cell lines

The knockout cell lines were generated via CRISPR-Cas9 system. sgRNAs targeting the indicated genes were designed by the online tool (https://chopchop.cbu.uib.no/). The corresponding DNA was synthesized and cloned into lenti-CRISPR-v2 puro, lenti-CRISPR-v2 BSD knockout vector. 17Cl-1 cells were infected with lentivirus collected from HEK293T cells and incubated for 48 h with 8 μg/mL polybrene. Transduced cells were selected with 2 μg/mL puromycin and 30 μg/mL Blasticidin (Cat. No. ST551, Cat. No. ST018) for 2 weeks. Single cell colonies were verified by immunoblotting analysis and sequencing of the PCR products. The *hgs* targeting sequence was TCTGCGACCTGATCCGTCAG.

### Co-IP and AP-MS

Co-IP and IB analysis were performed as described. In brief, for endogenous co-IP, 17Cl-1 cells (2 × 10^6^ cells) were seeded into a 10 cm dish and infected with MHV (MOI = 1). After 20 h, the total cells were harvested and lysed in 1ml ice-cold IP-lysis buffer (50 mM Tris-HCl pH 7.4, 5 mM EDTA, 40 mM β-sodium glycerophosphate, 30 mM NaF, 1 mM PMSF, 1 mM Na_3_VO_4_, 10% glycerol, 1.0% NP-40, and 150 mM NaCl) in the presence of a protease inhibitor cocktail (Roche) and phosphatase inhibitors. Cell lysates were precleaned and incubated with anti-HGS antibody and protein A/G agarose beads at 4 °C overnight. After washing five times, the immunoprecipitated complexes were sampled with SDS loading buffer and subjected to mass spectrometry analysis.

### IF

Cells infected with virus were fixed with 4% paraformaldehyde at room temperature for 15 min and permeabilized with 0.1% Triton X-100 and 3% BSA in PBS at room temperature for 1 h. Cells were then incubated with specific antibodies at 4 °C overnight, followed by an Alexa Fluor-labeled secondary antibody at room temperature in the dark for 2 h. Images of the cells were taken using a NIKON A1 inverted microscope or Carl Zeiss LSM 980. A list of antibodies used in this study is provided in the Supplementary Table S2.

### Live-cell imaging

HEK293T cells were seeded on a glass-bottom dish (Cat.No. D35-14-1-N). Cells were transfected with BFP-N, mCherry-M and GFP-HGS expressing plasmids using Lipofectamine 3000 Transfection Reagent (Cat. No. L3000001) for 24 h. Live-cell imaging was performed using Carl Zeiss Elyra 7 every 5 s. During live-cell imaging, the dish was maintained in the incubation conditions at 37 °C and 5% CO_2_.

### APEX2-based TEM

HEK-293T cells were transfected with a pEGFP vector expressing hHGS-APEX2, underwent fluorescence-activated cell sorting and were grown on ACLAR films. The cells were rinsed with phosphate-buffered saline (PBS), fixed with 2.5% glutaraldehyde in PBS for 1 h and then rinsed 3 times with PBS. The fixed cells were treated with 1.5 mg/mL glycine in PBS for 5 min to inactivate residual aldehyde and then rinsed 3 times with PBS. The cells were then submerged in 0.5 mg/mL 3,3′-diaminobenzidine (DAB) in PBS and kept in the dark for 40 min. Then aqueous hydrogen peroxide was added directly to the DAB solution so that a DAB solution supplemented with 10 μmol/mL H_2_O_2_ continued to treat the cells for another 15 min. The cells were rinsed 2 times with PBS and 2 times with double distilled water, before they underwent a standard sample preparation process as described above. Electron micrographs were acquired with an FEI Tecnai Spirit 120 kV transmission electron microscope operating at 100 kV accelerating voltage.

### Cryo-ET and cryo-CLIEM

Quantifoil R2/1 gold EM grids (200 mesh with holey carbon film of 2 μm hole size and 1 μm spacing) were plasma cleaned with O_2_ and H_2_ for 30 s in a glow discharge device (Gatan, Plasma Cleaner) and sterilized with UV light for 20 min in a biosafety cabinet. 17Cl-1 cells were seeded on EM grids in a 35 mm dish at a density of 1 × 10^5^ cells per dish and incubated at 37 °C overnight. Cells were transfected with plasmids (GFP-HGS) using Lipofectamine 3000 Transfection Reagent (Cat.No. L3000001). 24 h after transfection, samples were plunge-frozen in liquid nitrogen by using EM GP2 Automatic Plunge Freezer (Leica). Grids were blotted from single side (opposite to cell side) with 6 s blotting, chamber conditions of 25 °C and 85% humidity, and stored in liquid nitrogen. In the virus-infected samples, MHV viruses were added for infection after 14 h transfection. After 10 h infection, samples were plunge-frozen and stores in liquid nitrogen.

Vitrified cells were transferred to a cryo-correlated light, ion and EM (cryo-CLIEM) system that integrates a 3D multicolor confocal microscope into a dual-beam focused ion beam scanning electron microscope (FIB-SEM). Before FIB milling, 3D LM imaging was performed using an integrated confocal microscope and custom-made software. Tilt the images to same direction as the FIB milling, and determine the milling area according to the fluorescent signal. 500 pA currents were used for rough milling and lower current 50 pA for the polishing step to get the final lamella at 150 nm thickness. With no virus infection, a total of 7 lamella were prepared. In the virus-infected samples, a total of 8 cryo-lamella were prepared. The lamellas were imaged on a 300 kV Titan Krios electron microscope (Thermo Fisher Scientific) equipped with a field mission gun, a direct electron detector (Gatan, K2 camera) and an energy filter (Gatan). According to the fluorescence position, tilt-series were collected from −60° to +42° at 3° increments using SerialEM software in counting mode, with total accumulated dose of 100 e−/Å2, defocus of −6 μm, the energy filter of 20 eV and the final pixel size of 4.29 Å. Dose-fractioned images were aligned by MotionCor2. The corrected tilt-series were aligned with AreTomo and reconstructed by WBP implemented in the IMOD software package. The tomograms were segmented with EMAN2, before being polished and visualized with UCSF Chimera.

### Sample Preparation for FIB-SEM

17Cl-1 cells grown in 60mm dish were infected with MHV at MOI = 1. At 20 hpi the cells were rinsed with PBS and then fixed with 2.5% glutaraldehyde in PBS overnight. Extra post-staining steps were performed (the so-called OTO post-staining). Cells were post-fixed with osmium-ferricyanide (1% OsO_4_, 1.5% potassium ferrocyride) for 1 h at 4 °C in the dark. Further processing was done in the microwave. Cells were rinsed five times in dH_2_O for 15 min each and treated with 1% thiocarbohydrazide in dH_2_O for 20 min. Cells were rinsed three times with dH_2_O for 1 min each and stained four times with 1% OsO_4_ in dH_2_O for 1 h at 4 °C in the dark. Cells were rinsed three times with dH_2_O for 1 min, stained with 1% Uranyl acetate in dH_2_O for 1 h. The samples underwent a dehydration process in which the solution submerging the samples was sequentially replaced by 30%, 50%, 70%, 80%, 90%, 100% and 100% ethanol solutions, each grade submerging the sample for 10 min. Cells were infiltrated in Durcupan resin with increasing percentages of this resin in ethanol (25%, 50%, 75%) for 2 h, 8h and overnight in the microwave. Cells were embedded in a thin layer of Durcupan resin covered with a coverslip and polymerized overnight at 60 °C. After one day the resin slab was detached from the coverslips, polymerized at 45 °C for 12 h and at 60 °C for 48 h. The polymerized samples were mounted on SEM stubs.

### FIB-SEM

The resin-embedded block was pre-trimmed on the microtome to expose the sample surface, and then attached onto a 45° SEM stub. The entire sample was sputtered with gold to increase conductivity.

FIB-SEM image acquisition was performed using a dual beam scanning electron microscope (Aquilos2 Cryo-FIB, Thermo Fisher) with Auto Slice&View 10 Software (Thermo Fisher). Protective patch of platinum (∼3 μm thick)was deposited on the area of interest using ion beam deposition at 16 keV, 3.6 nA. A U-shaped trench was milled around the area of interest at 65 nA in order to expose the lamellar surface for the electron beam. The imaging surface was exposed by FIB milling of 50 μm deep trench at 30 kV, 3 nA in 7 °stage tilt. The slice thickness was set to 10 nm. The polish surface was imaged with the electron beam at 2 kV, 0.2 nA, 5 nm/pixel, 2 μs/pixel dwell time, 45° stage tilt, using the in-lens BSE detector (T1) in Optiplan mode. The automated ASV project run produced a 3D volume images at final voxel size of 5 × 5 × 10 nm³.

Alignment of image slices was performed by a workflow based on sequential linear registration. Pre-alignment was achieved by rigid transformation to compensate for translational and rotational deviations between consecutive slices. Subsequent alignment was refined using scale-invariant feature transform (SIFT)-based feature matching to ensure precise correspondence of local structural details. The displacements obtained from SIFT alignment were smoothed along the z-direction to enhance robustness and continuity, yielding the final aligned datasets.

Segmentation of DMVs, endomembrane system, multivesicular body, autophagolysosome, mitochondria and nucleus were performed in a semi- and fully automated manner in an original workflow that will be detailed elsewhere. The semi-automated workflow, based on machine learning-assisted segmentation using Arivis, was trained on manually annotated datasets to generate accurate ground truth for automatic segmentation. Post-processing, three-dimensional reconstruction, and rendering of the segmented organelles were subsequently performed within Imaris.

### Immuno-EM

17Cl-1 cells were cultured on ACLAR films and infected with MHV at MOI = 1. At 20 hpi, the cells were washed with PBS and chemically fixed using 4% PFA and 0.5% glutaraldehyde in PBS for 1 h. The fixed cells were then washed three times with PBS for 5 min each. The samples were subsequently treated with a blocking/permeabilization buffer (10% goat serum and 0.1% saponin in PBS) for 1 h. After three washes with 5% goat serum in PBS for 3 min each, the samples were treated with either (1) a dilution buffer (5% goat serum in PBS) or (2) the primary antibody (HGS antibody, 1:100 dilution in the dilution buffer) and incubated overnight at 4°C. The samples were then washed three times with 1% goat serum in PBS for 5 min each, before being incubated with the Nanogold secondary antibody (#2004; Nanoprobes, 1:50 dilution in the dilution buffer) for 1 h. Following this, the samples were washed three times with PBS and further fixed with 1% glutaraldehyde in PBS for 1 h. Any residual aldehyde was removed by three washes with 50 mM glycine in PBS for 5 min each. The samples were then washed three times with 1% bovine serum albumin in PBS for 5 min each and rinsed three times with ddH_2_O for 5 min each. Nanogold particles were enlarged using the Nanoprobes GoldEnhance EM Plus solutions according to the manufacturer’s instructions. The samples were rinsed three times with ddH_2_O for 5 min each before undergoing the standard TEM sample preparation protocol.

### Quantification and statistical analysis

The data were obtained from at least three independent experiments (n ≥ 3), and the representative data are presented as the mean ± SD as indicated and were corrected for multiple comparisons. ANOVA and Student’s t tests were used to analyze differences in mean values. GraphPad Prism 8.0.1 software was used to calculate the P-values, and significance is depicted with asterisks as follows: *P ≤ 0.05, **P ≤ 0.005, ***P ≤ 0.0005, ****P ≤ 0.0001.

## Supplementary Figure legends

**Supplementary Figure 1.**
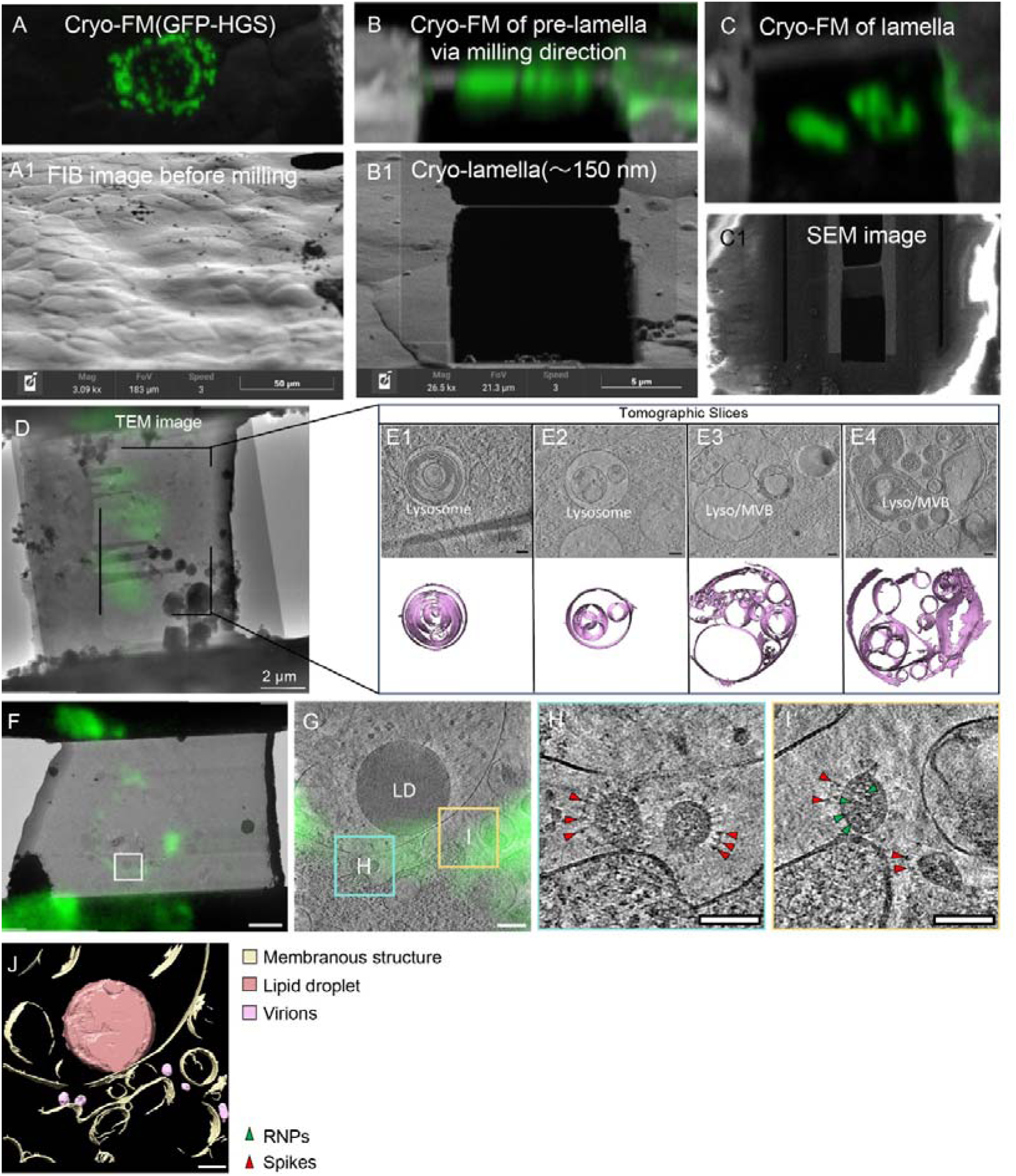
Cryo-CLEM analysis of HGS-GFP in 17Cl-1 cells and WIV1-infected Vero E6 cells. (A) Fluorescent GFP-HGS signals in vitrified 17Cl-1 cells. (A1) FIB image of same area. (B) Cryo-fluorescence microscopy (FM) image of lamella (∼300 nm) during FIB milling. (B1) Final lamella (∼150 nm), FIB view. (C) Cryo-FM image of final lamella (∼150 nm), top view. (C1) SEM image of final lamella. (D) Cryo-EM image colocalized with GFP-HGS puncta. (E1-E4) Gallery of tomographic slices and segmentations of vesicular organelles guided by GFP-HGS. Scale bars, 100 nm (E). (F-J) Cryo-CLEM analysis of HGS-GFP-expressing Vero E6 cells infected with WIV1. (F) HGS-GFP fluorescence (green) overlaid on low-magnification TEM map. Boxed area indicates cryo-ET acquisition site. (G) Tomographic slice from boxed area in F. LD: lipid droplet. (H) Enlarged view of cyan-boxed area in G, containing two WIV1 virions. (I) View of yellow-boxed area in G (different z-slice), containing two WIV1 virions. (J) 3D segmentation of the tomogram shown in G. Red arrowheads: spikes. Green arrowheads: RNPs. Scale bars: 2 μm (F), 200 nm (G and J), 100 nm (H and I).

**Supplementary Figure 2.**
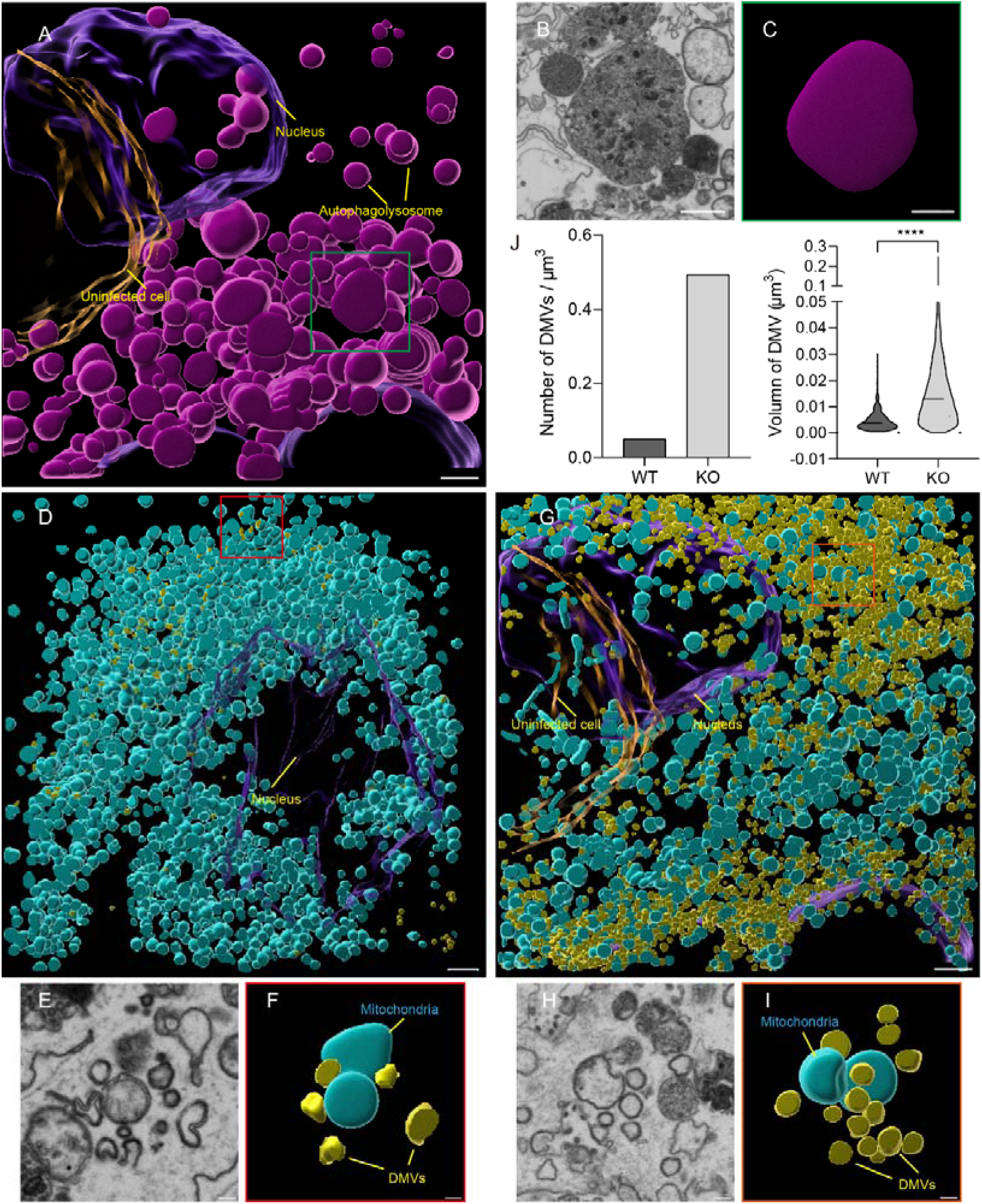
Volume EM reveals that HGS deficiency induces the formation of autophagolysosome, increases the density and volume of DMVs and impairs the mitochondria. (A-C) 3D rendering of Autophagolysosome in *Hgs*-KO cells. Scale bars, 2 μm in (A). (B) Zoom-in of the area indicated with rectangle in A. Scale bars, 1 μm. (C) same region as in (B) with superimposed rendering of autophagolysosome. (D-I) 3D rendering of DMV and mitochondria in WT (D-F) and *Hgs*-KO (G-I) cells. (E and F) Zoom-in of the area indicated with rectangle in D. (F) same region as in (E) with superimposed rendering of DMV and mitochondria in WT cells. (H and I) Zoom-in of the area indicated with rectangle in D. (I) same region as in (H) with superimposed rendering of DMV and mitochondria in *Hgs*-KO cells. Scale bars: 2 μm (D and G), 500 nm (E, F, H and I). (J) Quantification of density and volume of DMVs in WT and *Hgs*-KO cells. 394 and 3471 DMVs were selected randomly and calculated in WT and *Hgs*-KO cells, respectively. Data are the mean ± SD. Significance testing for (J) was performed with an unpaired t test. *P ≤ 0.05, **P ≤ 0.005, ***P ≤ 0.0005, ****P ≤ 0.0001, ns, no significance.

**Supplementary Figure 3.**
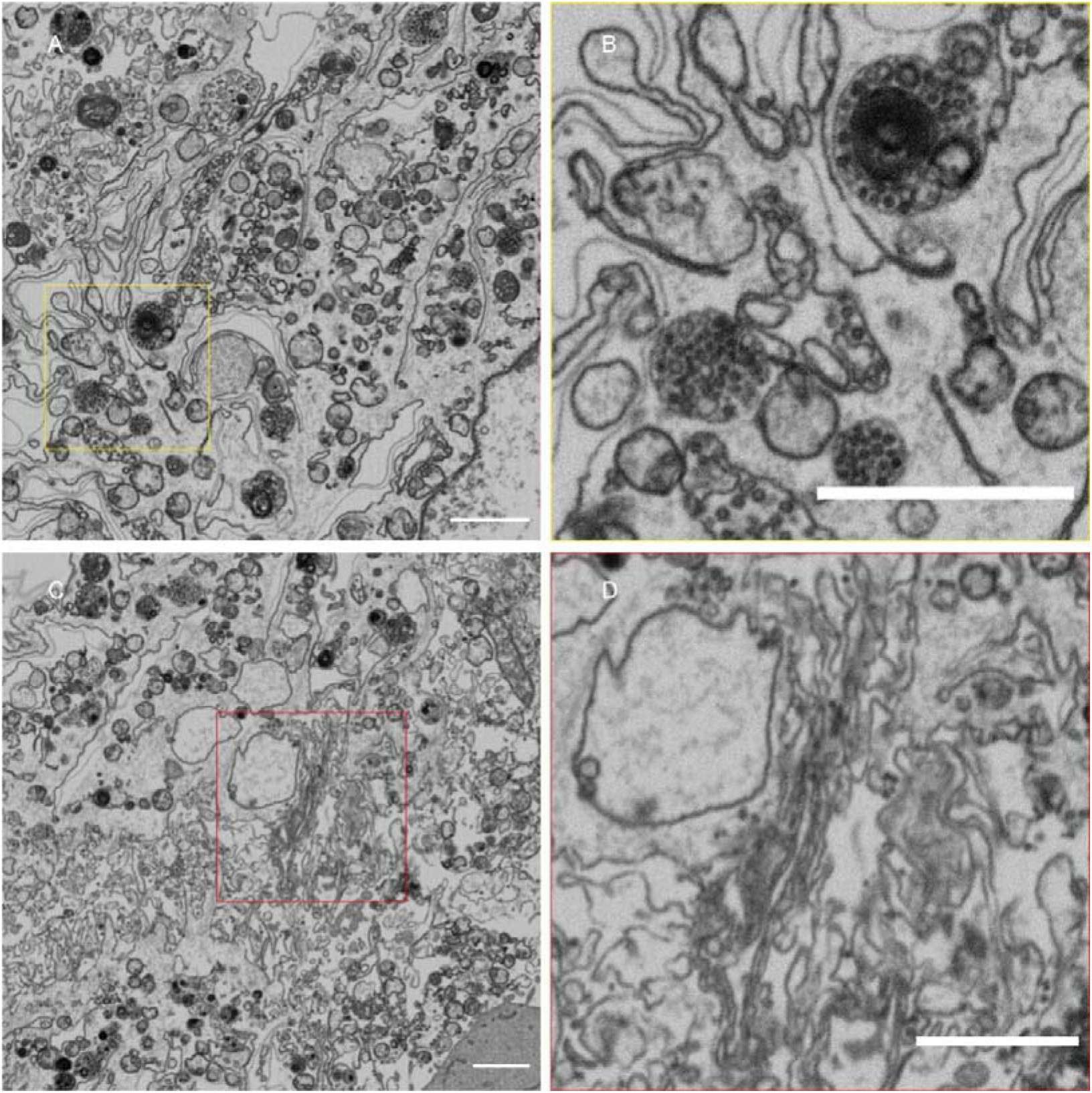
FIB-SEM analysis of MHV-infected 17Cl-1 cells reveals lysosomes and Golgi stacks. (A, C) Two different slices through the cell volume EM, showing lysosomes and Golgi stacks, respectively. (B, D) Zoom-in of the area indicated with rectangle in (A and C) showing lysosomes contain electron-dense luminal virions (B) and Golgi stacks adjacent to CVCs (D), respectively. Scale bars, 2 μm.

**Supplementary Table S1.** List of Cell lines.

|  |  |  |
| --- | --- | --- |
| Human: HEK293T | ATCC | Cat#<br>CRL-3216 |
| Human: Huh7 | Cas9X<sup>TM</sup> | Cat#<br>TCH-C217 |
| Human: Huh7.5.1 | Gifted by<br>Xiancai Ma | N/A |
| Mouse: 17Cl-1 | Gifted by<br>Hongyu Deng | N/A |
| Human: HRT-18 | Gifted by<br>Jincun Zhao | N/A |
| Monkey: LLC-MK2 | Gifted by<br>Jincun Zhao | N/A |
| African green monkey: Vero E6 | Gifted by<br>Xiancai Ma | N/A |
| Mouse: 17Cl-1 *Hgs*<sup>KO</sup> | This paper | N/A |

**Supplementary Table S2.**
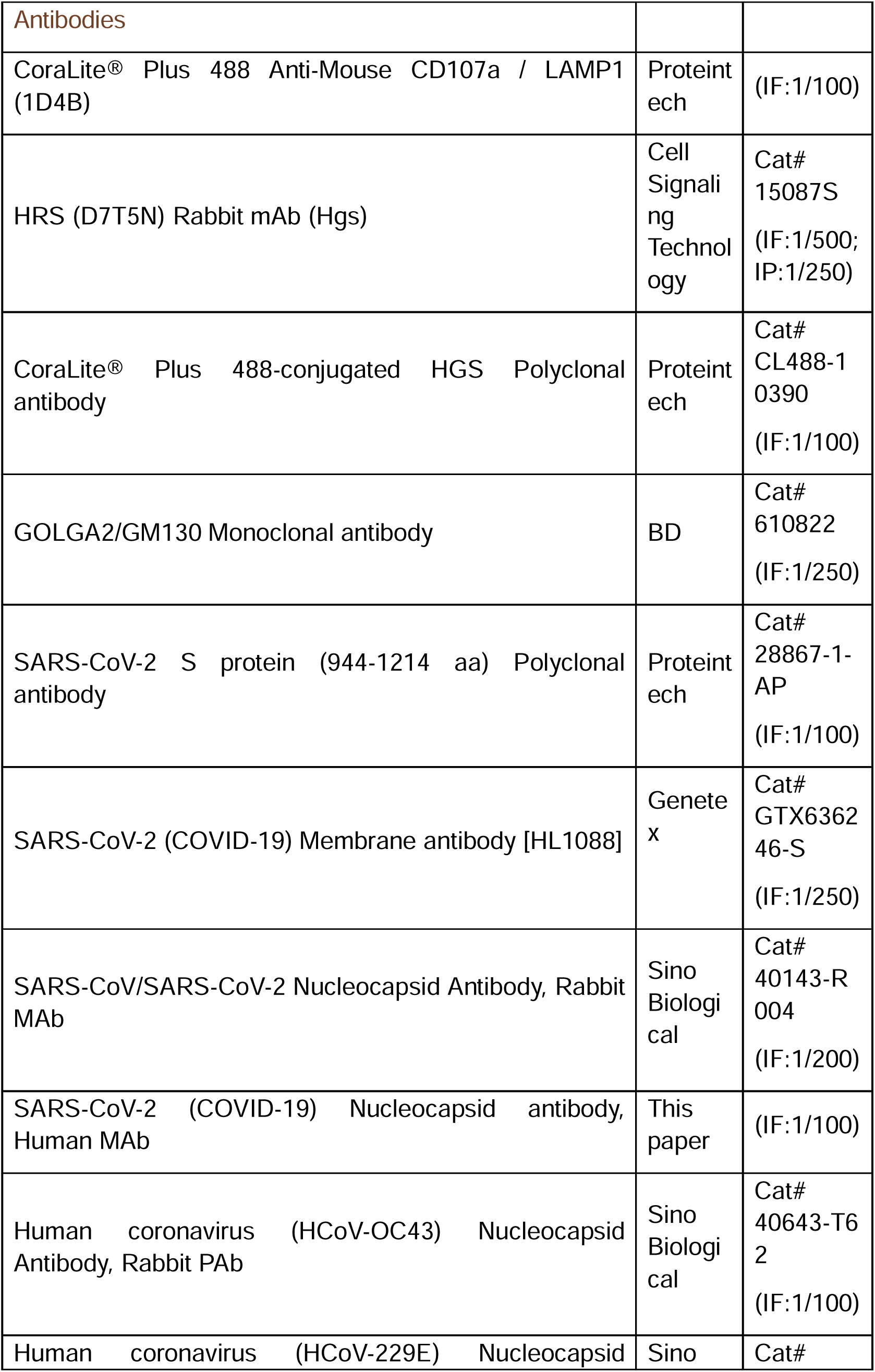
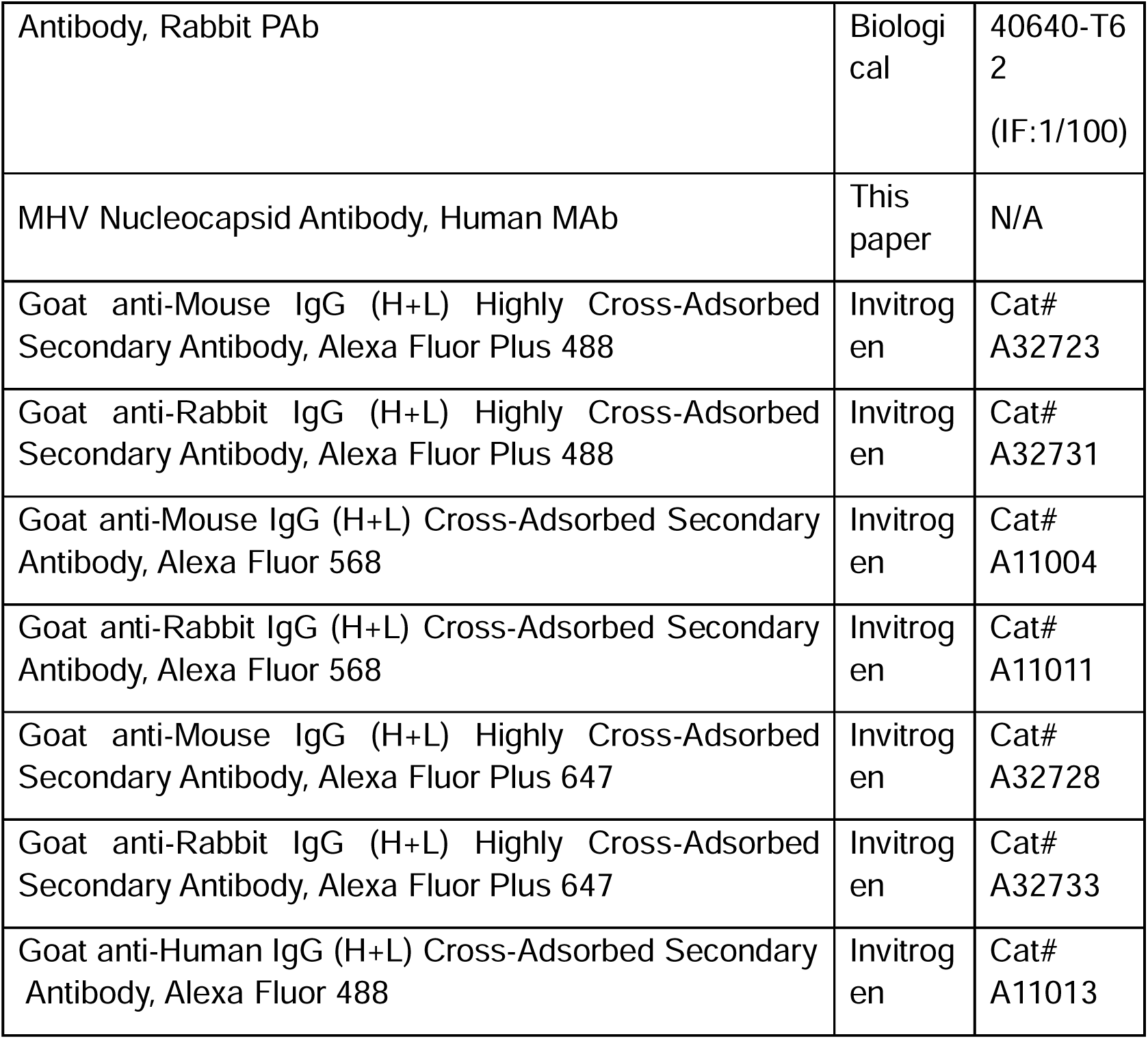
List of antibodies.

**Supplementary Table S3.** List of Chemicals.

|  |  |  |
| --- | --- | --- |
| Hoechst 33342 | Thermo<br>Fisher | Cat#<br>H1399 |
| Puromycin Dihydrochloride | Beyotime<br>Biotechnology | Cat#<br>ST551 |
| Blasticidin S HCl | Beyotime<br>Biotechnology | Cat#<br>ST018 |
| Polybrene (Hexadimethrine Bromide) | Beyotime<br>Biotechnology | Cat#<br>C0351 |
| Protein A/G Agarose Beads | Proteintech | Cat#<br>PR40025 |
| Lipofectamine™ 3000 Transfection Reagent | Thermo<br>Fisher | Cat#<br>L3000015 |
| TRIzol™ Reagent | Invitrogen | Cat#<br>15596018 |
| Fetal Bovine Serum, Premium Plus | Thermo<br>Fisher | Cat#<br>A5669701 |
| TrypLE™ Express Enzyme (1 × ), phenol red | Thermo<br>Fisher | Cat#<br>12605028 |
| Opti-MEM® I Reduced Serum Medium | Gibco | Cat#<br>31985070 |
| Penicillin-Streptomycin, Liquid (100 × ) | Invitrogen | Cat#<br>15140122 |
| EMS Aqueous Glutaraldehyde EM Grade 25% | Electron<br>Microscopy<br>Sciences | Cat#<br>16220 |

